# Dissecting aggregation and seeding dynamics of α-Syn polymorphs using the phasor approach to FLIM

**DOI:** 10.1101/2022.02.09.479740

**Authors:** Jessica Tittelmeier, Silke Druffel-Augustin, Ania Alik, Ronald Melki, Carmen Nussbaum-Krammer

## Abstract

Synucleinopathies are a heterogenous group of neurodegenerative diseases characterized by the progressive accumulation of pathological α-synuclein (α-Syn). The importance of structural polymorphism of α-Syn assemblies for distinct synucleinopathies and their progression is increasingly recognized. However, the underlying mechanisms are poorly understood. Here we use fluorescence lifetime imaging microscopy (FLIM) to investigate seeded aggregation of α-Syn in a biosensor cell line. We show that conformationally distinct α-Syn polymorphs exhibit characteristic fluorescence lifetimes. FLIM further revealed that α-Syn polymorphs were differentially processed by cellular clearance pathways, yielding fibrillar species with increased seeding capacity. Thus, FLIM is not only a powerful tool to distinguish different amyloid structures, but also to monitor the dynamic process of amyloid remodeling by the cellular environment. Our data suggest that the accumulation of highly seeding competent degradation products for particular polymorphs may account for accelerated disease progression in some patients.

## Introduction

The accumulation and prion-like propagation of amyloid deposits is a hallmark of many neurodegenerative diseases. Aggregated SNCA/α-synuclein (α-Syn) is associated with diseases termed synucleinopathies, including Parkinson’s disease and multiple system atrophy. Despite their common cause, synucleinopathies are highly heterogeneous. These different pathologies may be due to the fact that α-Syn forms conformationally distinct amyloid assemblies (Bousset et al., 2013; Hoppe et al., 2021; Shahnawaz et al., 2020; Strohäker et al., 2019; Van der Perren et al., 2020). Indeed, distinct α-Syn polymorphs have been shown to differ in their aggregation and propagation propensities and levels of toxicity *in vivo* (Bousset et al., 2013; Hoppe et al., 2021; Van der Perren et al., 2020). Moreover, the cellular environment seems to play a crucial role (Peng et al., 2018).

To decipher the progressive accumulation and spreading of disease-related proteins, the underlying mechanisms need to be studied in more detail in the cellular context. Biochemical assays have been widely used to analyze aggregation, but they are often too indirect and require the extraction of aggregated proteins from whole cell or animal lysates, which could lead to artifacts. Imaging techniques can be used to study aggregation *in situ* without the need for cellular lysis.

Methods such as fluorescence recovery after photobleaching (FRAP) and fluorescence loss in photobleaching (FLIP) have been used in an attempt to improve aggregate characterization. Unfortunately, these techniques only distinguish between mobile and immobile aggregation states, but clearly there are more aspects to an aggregate, such as the specific packing of the individual monomers in an amyloid fiber that remain invisible.

Aggregation into an amyloid fiber leads to quenching of an attached fluorophore due to compaction and crowding, thus reducing its fluorescence lifetime (Chen et al., 2016; Schierle et al., 2011). This can be measured using fluorescence lifetime imaging microscopy (FLIM) (Chen et al., 2016; Schierle et al., 2011). Therefore, FLIM is increasingly used to investigate aggregation processes in various *in vivo* models (Esbjörner et al., 2014; Laine et al., 2019; Sandhof et al., 2020). However, interpretation of fluorescence lifetimes becomes challenging when resolving lifetimes requires tedious multiexponential decay curve fitting, which is often the case with biological samples (Verveer, Squire, & Bastiaens, 2000). Here we use the phasor plot method to visualize the complex nature of aggregation in a fit-free manner. Each pixel of an intensity image is mapped to a point in the phasor plot corresponding to the mean of the measured fluorescence lifetime (Malacrida et al., 2021). Single exponential lifetimes and complex decays lie on the ‘‘universal circle” or within, respectively, with longer lifetimes located the closest to the (0,0) coordinate, while shorter lifetimes are shifted toward the right (Digman et al., 2008). This approach provides a convenient and fast global view of the fluorescence lifetime distribution, and unveils the high complexity of amyloid aggregation dynamics in the cellular environment.

In this study, we utilized the phasor approach to FLIM to monitor seeded aggregation of an α-Syn reporter construct in cell culture following the introduction of *in vitro* generated α-Syn fibrils. We show that different α-Syn polymorphs display distinct signatures in FLIM, which reflects their distinct folds as well as individual interactions with the cellular environment. FLIM also revealed that the seeds are getting processed by the cellular protein quality control network over time, as evidenced by a gradual increase in fluorescence lifetime. However, instead of rapid solubilization or degradation, α-Syn species with intermediate fluorescence lifetimes accumulated that preferentially localized to the seeded endogenous α-Syn aggregate. Although some polymorphs were differentially susceptible to certain protein clearance pathways, blocking their cellular processing reduced the formation of these intermediates as well as endogenous α-Syn aggregates. Thus, an incomplete cellular clearance of the amyloid seed accelerates seeded aggregation of native α-Syn, which could play a role in the progressive spreading of disease pathology with age. We further conclude that FLIM provides an additional window to features that remain hidden with conventional intensity-based imaging techniques.

## Results

### Monitoring seeded aggregation of α-Syn by phasor-FLIM

To evaluate the use of the phasor approach to FLIM in order to investigate seeded aggregation of α-Syn, HEK cells expressing A53T mutant α-Syn fused to yellow fluorescent protein (α-SynA53T-YFP) (Sanders et al., 2014) were treated with different *in vitro* assembled α-Syn fibers (Supplementary Fig. 1A). Under normal growth conditions, these cells display mainly a diffuse α-SynA53T-YFP signal (Fig. 1A). Addition of Fibrils resulted in the formation of distinct cytosolic foci in the fluorescence intensity image and a shift towards shorter α-SynA53T-YFP lifetimes in the corresponding phasor plot (Fig. 1A). In the phasor plots, a specific region of interest (ROI) can be selected to highlight pixels with that particular lifetime in the corresponding fluorescence intensity image (Ranjit et al., 2018). Selecting an ROI around shorter lifetimes (purple cursor) in the phasor plot reveals that these pixels colocalize with foci (Fig. 1B). Due to the reciprocity between intensity image and phasor plot, selecting an ROI around foci in the intensity image leads to the selective visualization of these pixels in the phasor plot, confirming that the lifetime of fluorophores within foci is shortened (Supplementary Fig. 1B). Hence, the seeded aggregation of endogenous α-SynA53T-YFP by the addition of exogenous α-Syn Fibrils leads to a shift in α-SynA53T-YFP lifetime distribution towards shorter lifetimes and the accumulation of short-lived protein species into distinct cytosolic foci.

**Figure 1.**
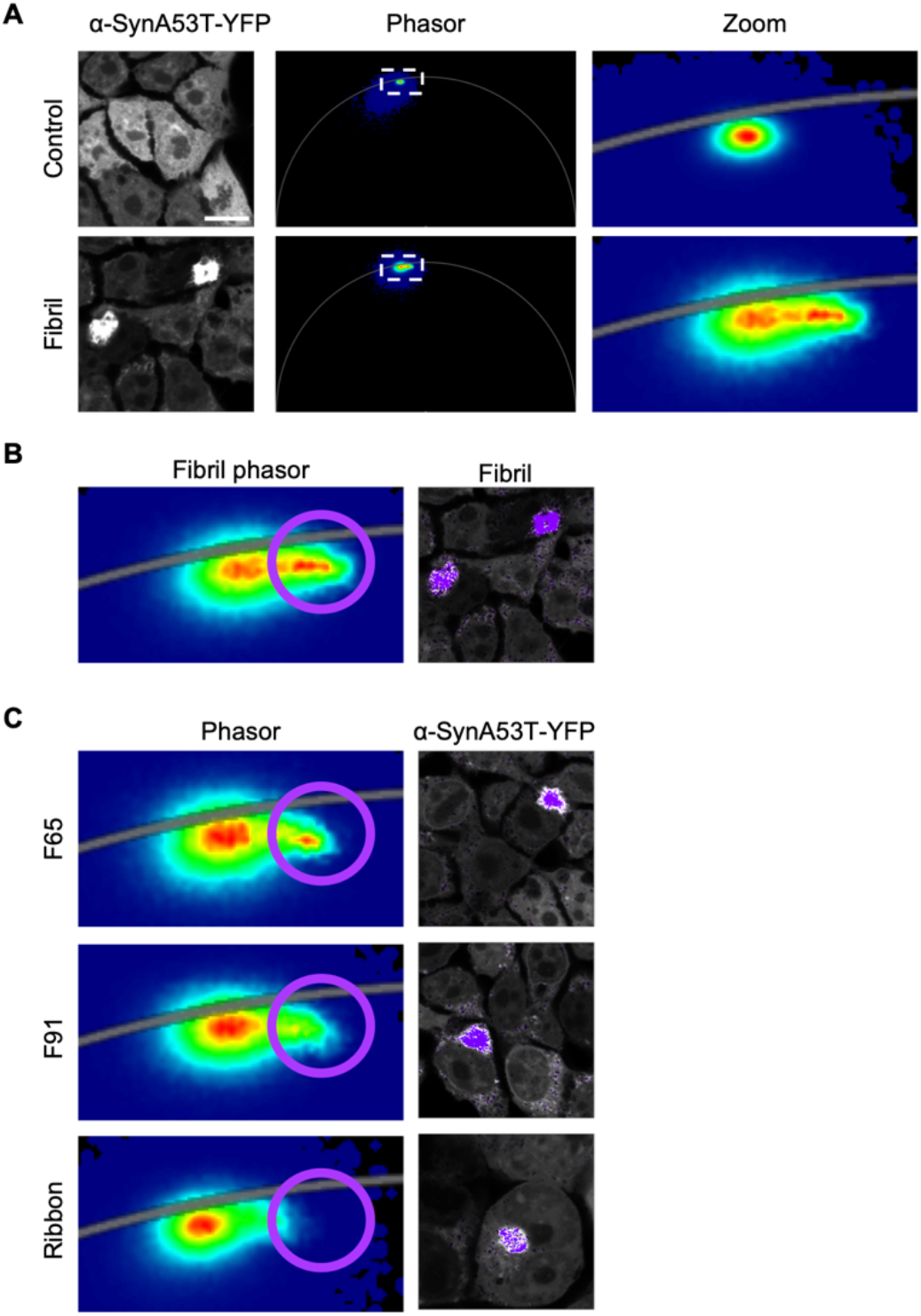
FLIM analysis of seeded aggregation of α-SynA53T-YFP using distinct α-Syn polymorphs. (A) Fluorescence intensity images of HEK cells expressing α-SynA53T-YFP seeded with indicated α-Syn fibers (left panel). Although most of the signal is from soluble α-SynA53T-YFP, the corresponding phasor plots of YFP fluorescence lifetimes show a shift in the distribution towards shorter lifetimes with seeded aggregation (middle and right panel). Scale bar, 10 μm. (B) Choosing a region of interest (ROI, purple) around shorter lifetimes in the phasor plot generates a pseudo colored image in which each pixel with a lifetime within the ROI is colored according to the color of the cursor in the phasor plot (purple cursor). The position of the cursor is arbitrary. The lifetime distribution of the non-seeded α-SynA53T-YFP in control cells was used as a reference to select the shorter α-SynA53T-YFP lifetimes that occur only in the seeded cells. Shorter lifetimes localize to distinct cytosolic foci. (C) Distinct α-Syn polymorphs are able to induce a shift in the α-SynA53T-YFP lifetime distribution towards shorter lifetimes that colocalize with foci.

α-Syn can form amyloid assemblies with distinct structures, so called conformational polymorphs, which exhibit distinct pathological characteristics *in vitro* and *in vivo* (Rey et al., 2019; Shrivastava et al., 2020). Based on the prion-like propagation the distinct conformations are often imposed on native α-Syn by templated seeding (Jones & Surewicz, 2005; Ke et al., 2020). Therefore, it was of interest to investigate whether the conformation of the fibrillar α-Syn seeds could have a measurable effect on the fluorescence lifetime of the seeded α-SynA53T-YFP aggregates. The addition of different α-Syn polymorphs (F65, F91 and Ribbons) led to the formation of α-SynA53T-YFP foci with shorter lifetimes similar to Fibrils (Fig. 1C, Supplementary Fig. 1C). However, although the different polymorphs had different seeding capacities (Supplementary Fig. 1D), no robust difference in lifetimes of α-SynA53T-YFP upon seeding with distinct α-Syn polymorphs could be observed.

### The different α-Syn polymorphs show a distinct distribution of fluorescence lifetime in phasor-FLIM

We wondered whether FLIM was sensitive enough to detect conformational differences between amyloid assemblies, and therefore sought to elucidate whether the seeds themselves had different fluorescence lifetimes. To test this, the distinct α-Syn polymorphs were labeled with Atto647 for direct visualization (Supplementary Fig. 2A). This not only allows imaging of the seed within a cell but also monitoring of the fluorescence lifetime of both the endogenous α-SynA53T-YFP and the fibrillar α-Syn seeds (α-Syn-Atto647). We observed that seeding with α-Syn-Atto647 did not cause major differences in the lifetime distribution of α-SynA53T-YFP compared to seeding with unlabeled seeds (Fig. 1, Supplementary Fig. 2B, C). In contrast, there were massive differences between the lifetimes of various α-Syn-Atto647 polymorphs (Fig. 2). Pseudocoloring of the lifetime distribution on the phasor map was used to highlight the location of shorter-lived species (blue) and green, yellow, and red species representing their progressively longer-lived fluorescence lifetimes in the corresponding fluorescence intensity image (Fig. 2A, B). Notably, Fibril, F65, and F91 polymorphs displayed a broad range of fluorescence lifetimes, ranging from around 0.2 ns to 1.8 ns (Fig. 2C, D), whereby F65 seeds formed species with the shortest fluorescence lifetime that do not appear with Fibrils and F91, while F91 formed species with a fluorescence lifetime of no less than 0.5 ns. Conversely, Ribbons did not exhibit such a wide distribution of fluorescence lifetimes, with the majority of seeds exhibiting lifetimes between 1.5 ns and 1.8 ns. Intriguingly, monomeric α-Syn displays also fluorescence lifetimes between 1.5 ns and 1.8 ns (Fig. 2E). This indicates that Ribbons are essentially invisible in FLIM and are incorrectly detected as monomeric α-Syn, although they are clearly fibrillar (Supplementary Fig. 2A), which led us to remove them from all subsequent analysis. Nevertheless, all visible α-Syn polymorphs exhibit a characteristic distribution of fluorescence lifetimes when exposed to cells, which is revealed by phasor-FLIM.

**Figure 2.**
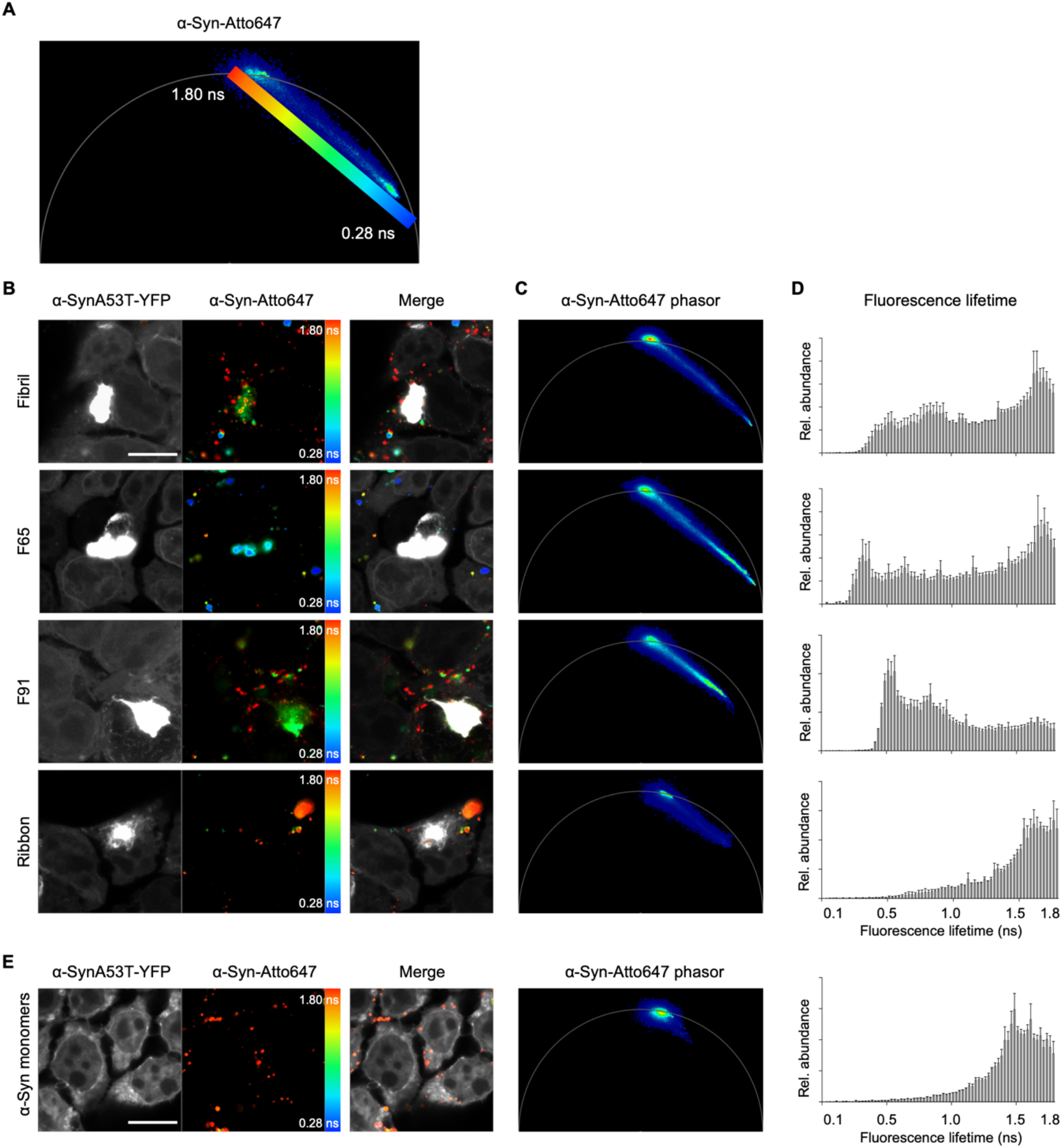
α-Syn polymorphs exhibit signature phasor fingerprints. (A) Pseudo coloring LUT from short fluorescence lifetime (0.28 ns) in blue to longer fluorescence lifetime (1.8 ns) in red on the phasor plot used in all α-Syn-Atto647 pseudo colored images. (B) Fluorescence intensity image of α-SynA53T-YFP and α-Syn-Atto647 seeds with pseudo colored from blue (0.28 ns) to red (1.8 ns) with the corresponding merged image. Scale bar, 10 μm. (C) Corresponding phasor plot of the fluorescence lifetimes of α-Syn-Atto647 seeds. (D) Histograms of α-Syn-Atto647 showing the relative abundance (rel. abundance) of pixels exhibiting a certain fluorescence lifetime (ranging from 0 to 1.8 ns), averaged over all acquired images (n = 15-20 for each polymorph, mean +SEM). Each α-Syn polymorph shows a signature fluorescence lifetime distribution. (E) Monomeric α-Syn displays a long fluorescence lifetime around 1.8 ns. Histograms of α-Syn-Atto647 monomers showing the relative abundance (rel. abundance) of pixels exhibiting a certain fluorescence lifetime (ranging from 0 to 1.8 ns), averaged over all acquired images (n = 15, mean +SEM). Scale bar, 10 μm.

### The cellular environment affects the lifetime distribution of α-Syn polymorphs

We wanted to further investigate the origin of the broad lifetime distributions of Fibril, F65, and F91 and asked whether this distribution is an intrinsic property of the particular polymorph. To test this, the fluorescence lifetime of α-Syn polymorphs was assessed directly. The initial seeds of Fibril, F65 and F91 each exhibited a specific short fluorescence lifetime with only a very narrow distribution (Fig. 3A, Supplementary Fig. 3A, B). Therefore, we hypothesized that the broad distribution of fluorescence lifetime (Fig. 2B-D) could be attributed to the intracellular environment. We asked whether it is caused by the interaction of the fluorophore Atto647 with YFP during the seeding process. To test this, fibrillar seeds were added to HEK cells not expressing α-SynA53T-YFP. The α-Syn polymorphs still exhibited their distinct broad distribution of fluorescence lifetimes when introduced into HEK cells (Supplementary Fig. 3C, D). Hence, the signature fluorescence lifetime fingerprint of the polymorphs in cells is not dependent on their interaction with α-SynA53T-YFP.

**Figure 3.**
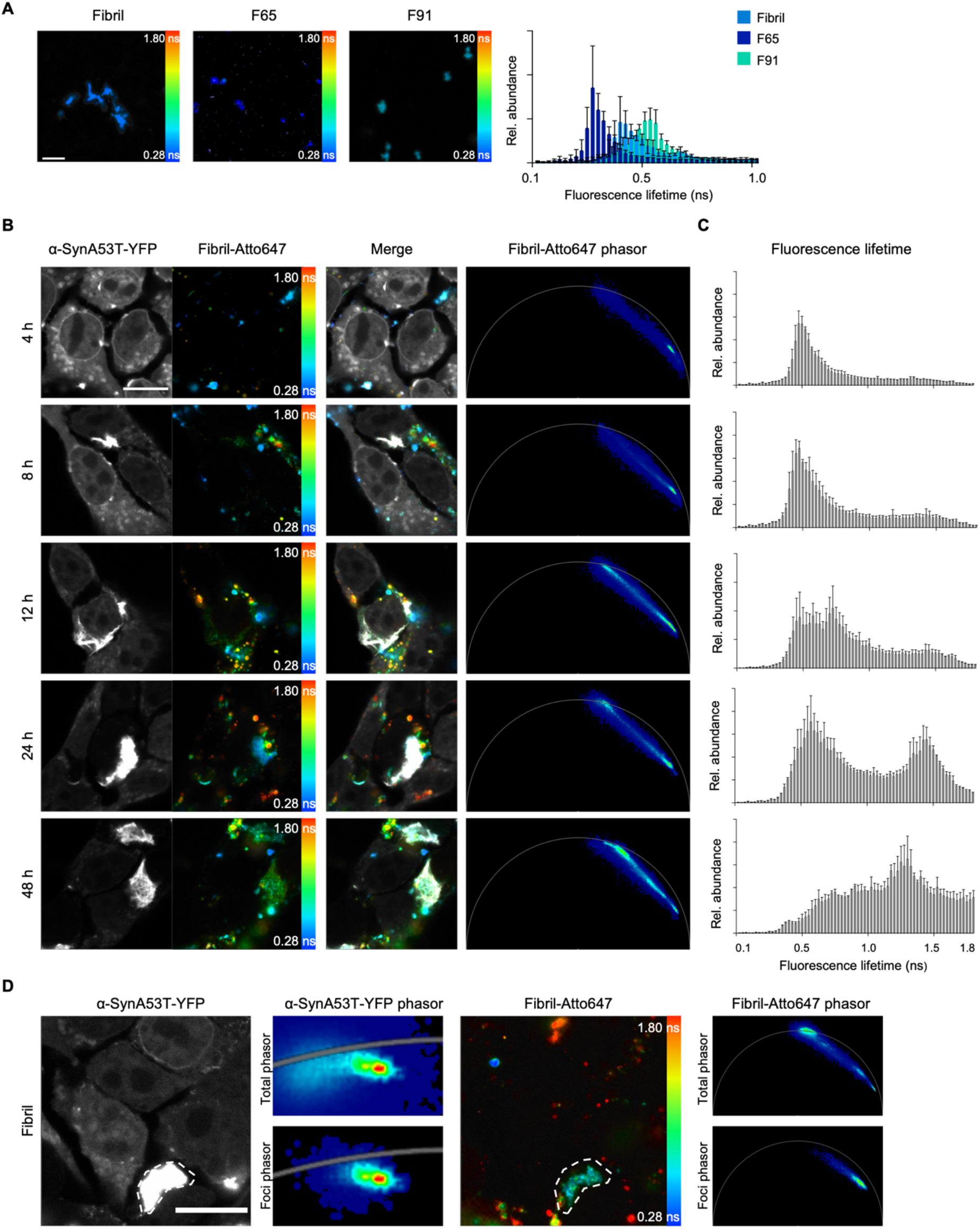
Kinetics of Fibril seed processing. (A) Fluorescence intensity image of α-Syn-Atto647 seeds pseudo colored from blue (0.28 ns) to red (1.8 ns) to illustrate short (blue) and long (red) fluorescence lifetimes when not added to cells with histograms of Fibril, F65 and F91 seeds showing the relative abundance (rel. abundance) of pixels with a certain fluorescence lifetime, averaged over all acquired images (n = 15 for each polymorph, mean + SEM). Each α-Syn polymorph seed shows a signature fluorescence lifetime distribution. Scale bar, 5 μm. (B) Fluorescence intensity images of α-SynA53T-YFP and α-Syn-Atto647 seeds pseudo colored with the merged images and the corresponding phasor plots of the fluorescence lifetime of Fibril-Atto647 seeds at indicated timepoints after seeding. Scale bar, 10 μm. (C) Histograms of Fibril-Atto647 seeds showing the relative abundance (rel. abundance) of pixels with a certain fluorescence lifetime at each timepoint, averaged over all acquired images (n = 15 for each timepoint, mean + SEM). Fibril-Atto647 seeds with longer fluorescence lifetimes accumulate in a time-dependent manner. (D) The ROI drawn around α-SynA53T-YFP foci (dotted white line) contains Fibril-Atto647 seed species with an intermediate lifetime (light blue) compared to the overall distribution of lifetimes (from dark blue to red). Scale bar, 10 μm.

Since the broad distribution of fluorescence lifetimes seems to come from exposure to the cellular environment, we set out to investigate the kinetics behind α-Syn seeds developing longer fluorescence lifetimes. While after 4 hours the internalized Fibril-Atto647 seeds resemble the initial Fibril-Atto647 seeds not exposed to cells, there is a time dependent accumulation of species with increasingly longer fluorescence lifetimes (Fig. 3A-C, Supplementary Fig. 3A, B). At 12 h post seeding, the characteristic broad fluorescence lifetime distribution in the phasor can be observed for the first time. In accordance, the relative abundance of longer fluorescence lifetimes increases in the corresponding lifetime histograms (Fig. 3B, C). Between 12 h and 24 h after seeding, an increasing number of species with fluorescence lifetimes between 1.5 ns and 1.8 ns form. The latter are possibly monomers. This processing of the seeds over time is also seen for F65 and F91 (Supplementary Fig. 4A, B). We conclude that the large distribution of lifetimes observed for some α-Syn seeds is a result of cellular processing over time. Intriguingly, intermediate fluorescence lifetime species preferentially localize to α-SynA53T-YFP foci, rather than the likely unprocessed original seed with a very short fluorescence lifetime (Fig. 3D). We therefore speculated that there might be a correlation between the cellular processing of seeds and the seeding efficiency of α-Syn polymorphs.

### Processing of α-Syn fibrillar polymorphs by protein quality control systems aggravates seeding

We investigated whether cellular processing of a seed may give rise to a more seeding-competent species. There are several cellular pathways that might mediate this processing. In our approach, seeds are delivered directly into the cytosol of the cell by transfection. Three main protein quality control systems are responsible for cytosolic removal of protein aggregates. These are the autophagy-lysosomal system and the ubiquitin-proteasome system, both of which accomplish protein degradation, or chaperone-mediated disaggregation, which intends to dissolve aggregated proteins. We therefore tested the impact of these pathways on the processing of Fibrils. First, we targeted the lysosomal degradation pathway. Chloroquine (CQ) was added 12 h before seeding with Fibrils, to increase lysosomal pH and thus block autophagic flux (Mauthe et al., 2018). 12 h after seeding, the characteristic fingerprint of fluorescence lifetimes developed in the control, but to a lesser extent with lysosomal degradation blocked. A short-lived fluorescence lifetime was more prominently detected, reminiscent of the initially introduced seed (Fig. 4A, B, Fig. 3A). Remarkably, the formation of endogenous α-SynA53T-YFP foci also significantly decreased with inhibition of the autophagy-lysosomal system (Fig. 4G, Supplementary Fig. 5A). Thus, the broadening of the fluorescence lifetime distribution in FLIM appears to indeed reflect processing of Fibrils by lysosomal degradation. The results further suggest that this processing generates more seeding competent species.

**Figure 4.**
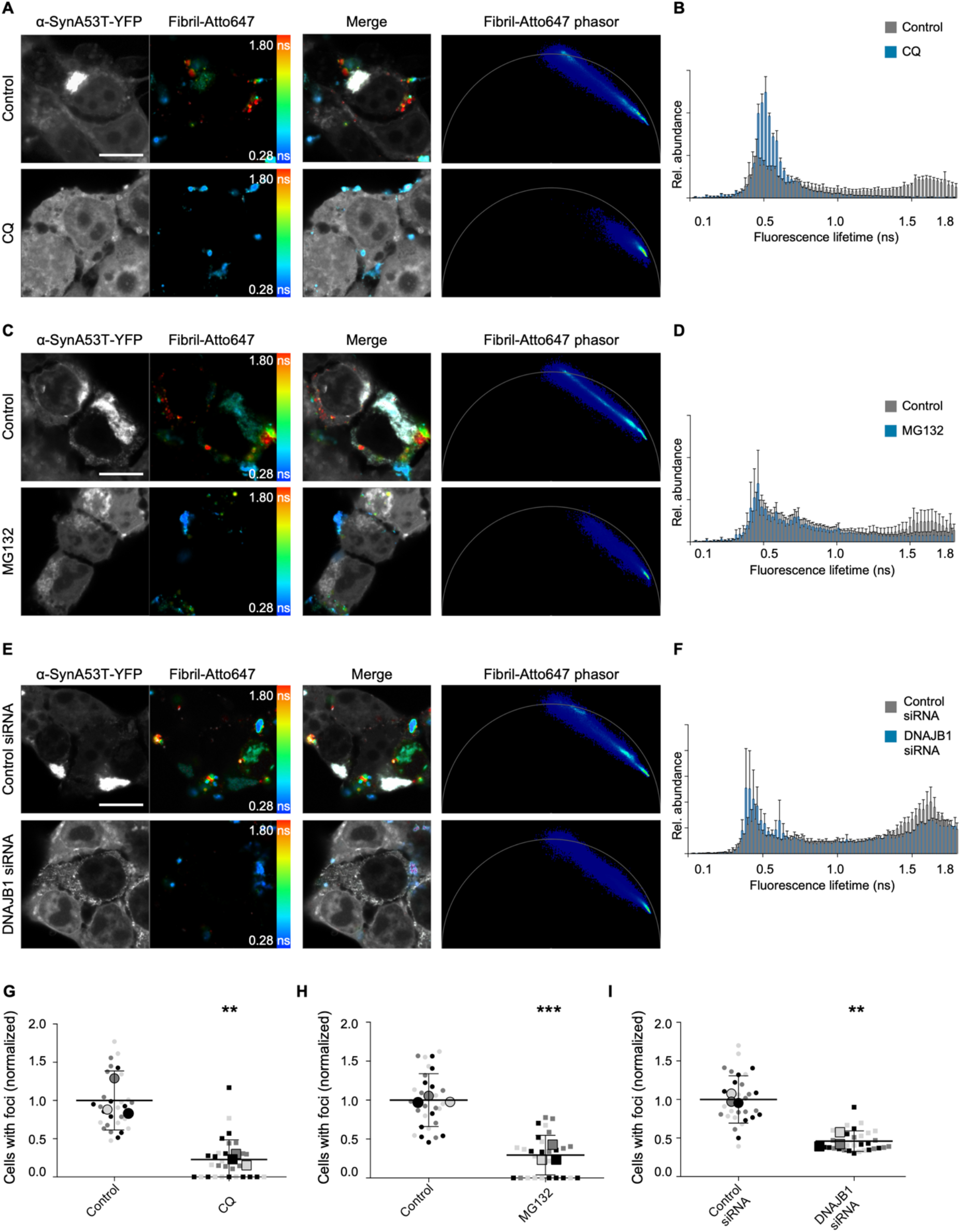
Protein clearance machinery augments α-Syn seeding. (A, C, E) Fluorescence intensity images of α-SynA53T-YFP and Fibril-Atto647 seeds pseudo colored from blue (0.28 ns) to red (1.8 ns) to illustrate short (blue) and long (red) fluorescence lifetimes with the merged images and the corresponding phasor plots of the fluorescence lifetime of Fibril-Atto647 seeds upon treatment with chloroquine (CQ) (A), MG132 (C), or DNAJB1 siRNA (E). Scale bar, 10 μm. (B, D, F) Histograms of Fibril-Atto647 seeds showing the relative abundance (rel. abundance) of pixels with a certain fluorescence lifetime (ranging from 0 to 1.8 ns), averaged over all acquired images (n = 15 for each condition, mean + SEM). (G-I) Quantification of cells with visible foci. For each biological replicate (n = 3) a mean was calculated; these three means were then averaged (horizontal bars) and used to calculate standard deviation (error bars), and p values. **p < 0.01, ***p < 0.001. Statistical analysis was done using an unpaired t test. Inhibition of the proteasome and autophagy, and silencing of the HSP110/HSP70 disaggregation machinery by DNAJB1 siRNA partially blocks the processing of the initial Fibril-Atto647 seed, which reduces α-Syn seeding.

To assess the role of the ubiquitin-proteasome system in the processing of Fibrils, we blocked the proteasome for 4 h using MG132 before the addition of Fibrils. Similar to the lysosomal degradation inhibition, this also partially impeded the development of the characteristic longer fluorescent lifetimes of Fibrils after 12h of seeding (Fig. 4C, D), and again was accompanied by a reduction in α-SynA53T-YFP foci formation in the cells treated with MG132 (Fig. 4H, Supplementary Fig. 5B). Hence, inhibiting the processing of Fibrils, either by blocking the autophagy-lysosomal system or the ubiquitin-proteasome system, reduced seeded aggregation of α-Syn.

The disassembly of amyloid aggregates by the HSP110/HSP70 disaggregation machinery has been shown to generate smaller seeding competent species (Nachman et al., 2020; Tittelmeier, Sandhof, et al., 2020). Therefore, we wondered whether targeting this disaggregation activity would also prevent the processing of Fibrils and impact α-Syn seeded aggregation. Disaggregation by the HSP110/HSP70 system depends on the substrate recognition and recruitment of HSP70 by the J-domain protein DNAJB1 (Gao et al., 2015; Wentink et al., 2020). Consequently, we targeted disaggregation by knocking down DNAJB1 prior to seeding (Supplementary Fig. 5C). DNAJB1 knockdown (KD) partially hampered the processing of the original Fibril seed into species with longer fluorescent lifetimes (Fig. 4E, F). Disaggregation of Fibrils also seemed to generate seeding competent species as there is a reduction in α-SynA53T-YFP foci formation in cells with reduced DNAJB1 levels (Fig. 4I). Of note, while DNAJB1 KD reduced foci formation in general, we noticed an increase in elongated foci as opposed to the typical spherical foci (Supplementary Fig. 5D zoom, 5E), suggesting that DNAJB1 may affect not only exogenously added seeds but also endogenous α-SynA53T-YFP aggregates. As a control, we targeted DNAJA2, which is involved in the prevention of aggregation rather than disaggregation (Faust et al., 2020; Gao et al., 2015). In accordance, DNAJA2 KD did not impact the fluorescence lifetime of Fibrils and actually slightly increased their seeding efficiency, in line with of the role in preventing α-Syn aggregation (Supplementary Fig. 5F-I).

When the treatments were repeated with F65 and F91 polymorphs, we found that the different protein clearance pathways could not process all α-Syn polymorphs to a similar extent as Fibrils. Whereas inhibition of the autophagy-lysosomal system blocked the processing of both F65 and F91 polymorphs (Fig. 5A, Supplementary Fig. 6A, Supplementary Fig. 7A), which was accompanied by reduced seeding efficiency (Fig. 5B, Supplementary Fig. 6B, Supplementary Fig. 7B), it differed for the ubiquitin-proteasome pathway. For F65, inhibition of proteasomal degradation diminished the generation of species with longer fluorescent lifetimes, while processing of F91 was only marginally affected by the MG132 treatment. In accordance, there was a reduction in formation of endogenous α-SynA53T-YFP foci only in the case of F65, but not in the case of F91 (Fig. 5C, D, Supplementary Fig. 6C, D, Supplementary Fig. 7C, D). This suggests that the proteasome may not be able to process the initial F91 seed, as is the case for F65 and Fibril.

**Figure 5.**
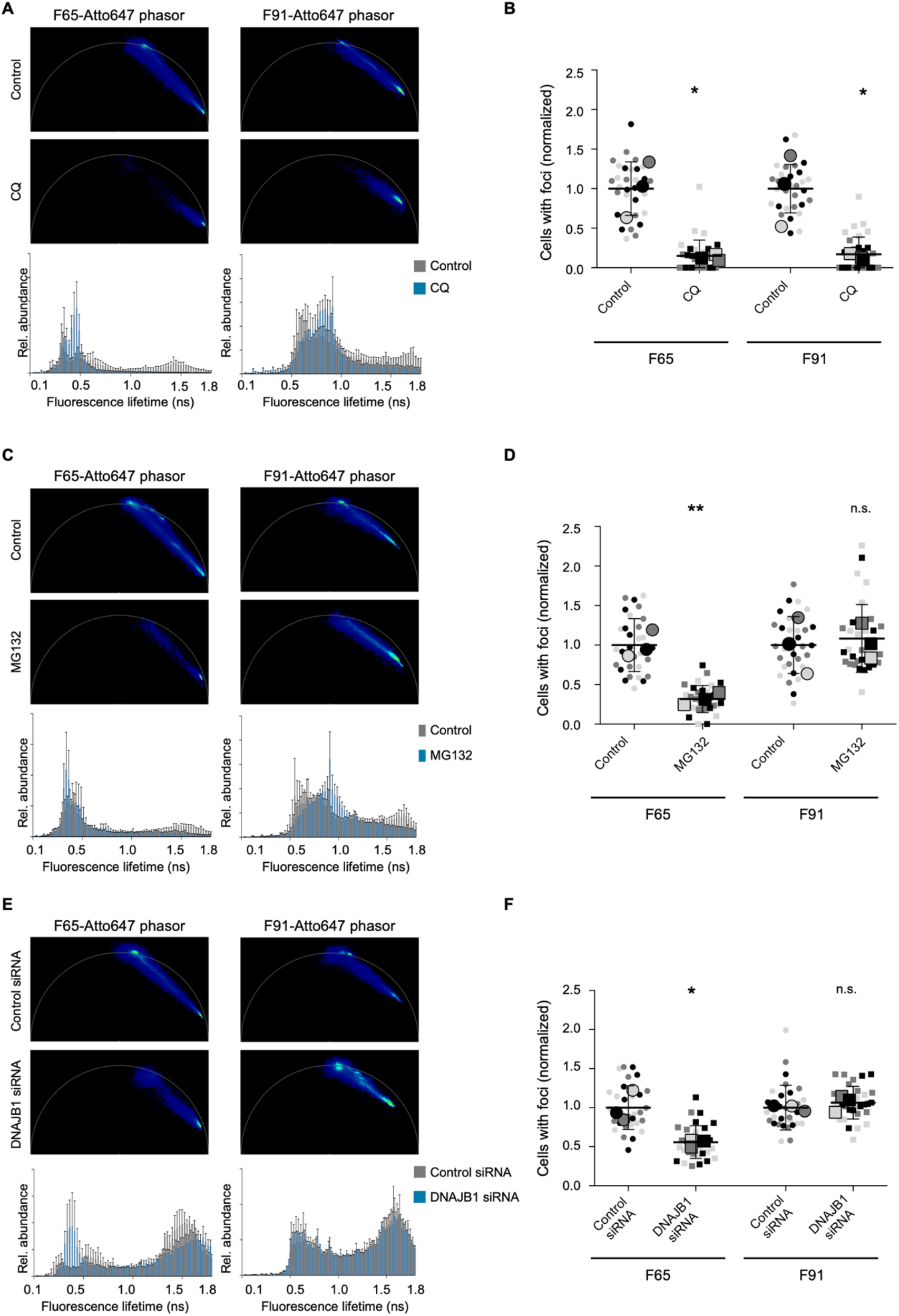
Differential processing of α-Syn polymorphs by the protein clearance pathways. (A, C, E) Phasor plots of the fluorescence lifetime of F65-Atto647 and F91-Atto647 seeds upon cells treatment with chloroquine (CQ) (A), MG132 (C), or DNAJB1 (E) siRNA. Histograms of F65-Atto647 and F91-Atto647 showing the relative abundance (rel. abundance) of pixels with a certain fluorescence lifetime (ranging from 0 to 1.8 ns), averaged over all acquired images (n = 15 for each condition, mean + SEM). (B, D, F) Quantification of cells with visible foci. For each biological replicate (n = 3) a mean was calculated; these three means were then averaged (horizontal bars) and used to calculate standard deviation (error bars), and p values. n.s. = not significant, *p < 0.05, **p < 0.01. Statistical analysis was done using an unpaired t test. Inhibition of autophagy (B) and the proteasome (D), and silencing of the HSP110/HSP70 disaggregation machinery by DNAJB1 siRNA (F) partially blocks the processing of F65-Atto647 seeds, which reduces α-Syn seeding. In contrast, only inhibition of autophagy partially blocks the processing of F91-Atto647 seeds.

Moreover, DNAJB1 KD partially suppressed the processing of the F65 seeds into species with longer fluorescent lifetimes resulting in a decrease in foci formation (Fig. 5E, F, Supplementary Fig. 6E, F). In contrast, DNAJB1 KD did not attenuate the processing of the F91 seeds and the formation of endogenous α-SynA53T-YFP foci (Fig. 5E, F, Supplementary Fig. 7E, F). Thus, F91 may not be recognized by DNAJB1 or may not be efficiently disaggregated by the HSP110/HSP70 system. Of note, DNAJA2 KD had no effect on the fluorescence lifetime nor the seeding efficiency for both F65 and F91 (Supplementary Fig. 6G-I, Supplementary Fig. 7G-I). These results suggest that the structural arrangement of α-Syn polymorphs impacts the processing ability of different protein clearance pathways.

Taken together, the broad distribution of fluorescence lifetimes appears to be attributable to the activity of multiple quality control pathways targeting aggregated proteins for degradation or disaggregation. Furthermore, the processing of α-Syn seeds by quality control pathways may produce fibrillar species with increased seeding capacity, which could accelerate the progressive spreading of pathogenic proteins.

## Discussion

The gradual accumulation and intercellular spreading of misfolded proteins is intimately linked to the development and propagation of neurodegenerative diseases. The ability of amyloidogenic proteins to form aggregates with different conformations might account for heterogeneity regarding pathology and progression (Kim et al., 2022; Melki, 2018). The cellular milieu also seems to have a crucial influence (Peng et al., 2018). Therefore, it is of great interest to investigate the role of amyloid conformation on the seeded propagation in the cellular context. Here, we describe a non-invasive fluorescence-based microscopy technique that enabled us to study the dynamics and molecular events of seeded α-Syn aggregation in a biosensor cell line to unprecedented detail. α-Syn polymorphs exhibit distinct signature fingerprints in FLIM, allowing the discrimination of conformational variants. Our data further establish a mechanistic link between amyloid processing by the cellular protein quality control machinery and seeding capacity and provide a possible basis for the correlation between the age-related decline of the proteostasis network and the progression rate of various synucleinopathies.

By using sequential multi-channel FLIM, we could visualize and distinguish both the endogenous aggregation of α-Syn and the fate of the exogenous α-Syn fibrillar polymorphs during the seeding process. Direct visualization of α-Syn fibrillar polymorphs labeled with Atto647 revealed that they exhibited differences in their fluorescence lifetimes. For example, F65 had the shortest initial fluorescence lifetime compared to any of the other polymorphs, while Ribbons showed relatively long fluorescence lifetimes despite their amyloid nature (Fig. 2, 3, Supplementary Fig. 3). These differences could be caused by different positions of the attached fluorophores given that labeling is achieved after the amyloids are formed. We indeed recently demonstrated that the exposure to the solvent of lysine residues within the different amyloid structures differs in distinct fibrillar polymorphs (Landureau et al., 2021). Fluorophores that are closer to the core of the fiber would be more quenched resulting in shorter lifetimes, while fluorophores that are closer to the N- or C-terminal end could move more freely resulting in longer lifetimes. Although this renders some polymorphs invisible in FLIM and precludes their analysis, it indirectly reflects differences in the amino acid residues that constitute the core of the fibers or the stacking of the individual monomers, and thus enables the discrimination of conformational variants. It would be interesting to compare the different α-Syn polymorphs in two-photon FLIM, which does not require labeling to examine whether this approach also allows their differentiation (Chung et al., 2021).

FLIM also allows detection of species that are formed during the seeding process. This technique revealed cellular processing of the α-Syn polymorphs over time as demonstrated by a gradual broadening of their fluorescence lifetimes, ranging from very short lifetime compositions, around 0.28 ns, to longer lifetimes at around 1.8 ns (Fig. 3, Supplementary Fig. 4) in the phasor plot.

We show that processing of fibrillar α-Syn appears to be mediated by multiple protein clearance pathways rather than a single one (Fig. 4 and 5). The HSP110/HSP70 disaggregation system attempts to dissolve fibers thereby releasing monomeric α-Syn and generating smaller fragments (Gao et al., 2015). A recent study showed that disaggregation of α-Syn fibrils *in vitro* leads to a shift of the fluorescence lifetime distribution towards longer lifetimes in FLIM (Chung et al., 2021). Accordingly, inhibition of the disaggregation activity in our cell culture model was reflected by a decrease in fiber lifetimes in FLIM accompanied by longer fluorescence lifetimes 24 h after seeding (Fig. 4F, 5E). The same pattern was seen by blocking proteasomal degradation, which is consistent with observations that the proteasome can also fragment and disassemble α-Syn fibrils (Cliffe et al., 2019; Sang et al., 2021). In addition, larger cytosolic α-Syn assemblies are known to be directed to the autophagy-lysosomal pathway for degradation (Webb et al., 2003), and overloading of this route could also lead to the release of smaller amyloidogenic particles (Flavin et al., 2017; Jiang et al., 2017; Marrero-Winkens et al., 2020). Furthermore, it is known that amyloid seeds are proteolytically processed by various cytosolic or lysosomal proteases after their uptake into the cell (Pieri et al., 2016). The cleavage of the N- and C-terminal fragments and attached fluorophores, might result in fewer intermolecular interactions of fluorophores. Whether and to what extent such proteolytic digestion contributes to the observed increase in fluorescence lifetimes, needs to be further investigated.

FLIM further revealed that structural differences in α-Syn polymorphs affected their susceptibility to different cellular aggregate clearance pathways, which could have implications in the different pathologies observed in synucleinopathies (Suzuki et al., 2020). Regardless of the particular pathway, the processing of α-Syn fibers by the proteostasis network generated species with high seeding capacity that enhanced aggregation of endogenous α-Syn (Fig. 4, 5, Supplementary Fig. 5, 6). Using the phasor approach to FLIM demonstrated that the endogenous α-Syn aggregates mainly formed around α-Syn seeds with intermediate fluorescence lifetimes, not the unprocessed α-Syn seeds with the shortest fluorescence lifetimes (Fig. 3D). Hence, the observed processing seems to promote seeded aggregation.

To the best of our knowledge, we show for the first time *in situ* that processing by the cellular proteostasis network does generate distinct α-Syn species with high seeding capacity. Our data offer an explanation for the conundrum that is increasingly observed with protein quality control pathways, as being generally protective and mediating the degradation of toxic protein aggregates, but sometimes having the opposite effect (Macošek et al., 2021; Tittelmeier et al., 2020). Indeed, there is mounting evidence that their activity can promote amyloid propagation under certain circumstances (Heiseke et al., 2009; Sang et al., 2021; Tittelmeier et al., 2020).

In our model system, the clearance pathways are likely overloaded by the addition of large amounts of exogenous seeds, probably leading to slower or incomplete disaggregation and/or degradation. This seems to favor the accumulation of intermediates that exhibit high seeding propensity (Franco et al., 2021; Marrero-Winkens et al., 2020). While this is an artificial system, the observation nevertheless has physiological relevance, as the capacity of the protein quality control system declines with age and to different extents in different tissues (Labbadia & Morimoto, 2015; Sabath et al., 2020). In particular, cellular degradation pathways such as the ubiquitin-proteasome system or the autophagy-lysosome pathway become increasingly overwhelmed during aging (Aman et al., 2021; Saez & Vilchez, 2014). This overload and associated accumulation of intermediate, highly seeding competent amyloid species could be a reason for the progressive spreading of disease pathology in neurodegenerative diseases.

## Materials and Methods

### *In vitro* aggregation of recombinant α-Syn

Human full length wild-type human α-Syn and human α-Syn lacking 30 C-terminal amino-acid residues (α-Syn 1-110) were generated in *E. coli* BL21 DE3 CodonPlus cells (Agilent Technologies) and purified as described (Ghee et al., 2005). To generate the fibrillar polymorphs Fibrils, Ribbons, F65 and F91, full-length monomeric α-Syn (250μM) was incubated in 20 mM Tris–HCl, pH 7.5, 150 mM KCl; in 5 mM Tris–HCl, pH 7.5; in 20mM MES pH 6.5 and in 20mM KPO4, 150mM KCl, respectively, at 37°C under continuous shaking in an Eppendorf Thermomixer set at 600 r.p.m for 7 days (Bousset et al., 2013; Makky et al., 2016). All assembly reactions were followed by withdrawing aliquots (10 μl) from the assembly reactions at different time intervals, mixing them with Thioflavin T (400μl, 10 μM final) and recording the fluorescence increase on a Cary Eclipse Fluorescence Spectrophotometer (Varian Medical Systems Inc.) using an excitation wavelength = 440 nm, an emission wavelength = 480 nm and excitation and emission slits set at 5 and 10 nm, respectively. The resulting α-Syn fibrillar polymorphs were assessed by Transmission Electron Microscopy (TEM) after adsorption of the fibrils onto carbon-coated 200 mesh grids and negative staining with 1% uranyl acetate using a Jeol 1400 transmission electron microscope. The fibrillar polymorphs were next labelled by addition of one molar equivalents of Atto-647 NHS-ester (#AD 647-35, ATTO-Tec GmbH) fluorophore in DMSO. The mix was incubated for 1 h at room temperature. The unreacted fluorophore was removed by two centrifugations at 15,000 *g* for 10 min and resuspensions of the pellets in PBS. The fibrillar polymorphs were next fragmented to an average length of 42-52 nm by sonication for 20 min in 2 mL Eppendorf tubes in a Vial Tweeter powered by an ultrasonic processor UIS250v (250 W, 2.4 kHz; Hielscher Ultrasonic) after assembly (Pieri, Madiona, & Melki, 2016). Monomeric α-Syn (250μM) in PBS was labeled by addition of one molar equivalent of Atto-647 NHS-ester for 2h on ice. The unreacted dye was removed by size exclusion chromatography on NAP-5 columns. Monomeric α-Syn-Atto-647 was aliquoted (10μl per 0.5 ml Eppendorf tube). Flash frozen in liquid nitrogen and stored at −80°C until use.

### Cell culture

Cells were cultured in DMEM containing high glucose, GlutaMAX Supplement, and pyruvate (Gibco), supplemented with 10% FBS (Gibco) and 1x Penicillin-Streptomycin (Gibco) at 37 °C and 5% CO_2_. Regular mycoplasma tests were performed (GATC Biotech). The HEK293T cell line expressing α-SynA53T-YFP was kindly provided by Marc Diamond, University of Southwestern Texas.

### Liposome-mediated seeded aggregation of cells and treatments

Cells were seeded with different preformed α-Syn fibrils. Cells were seeded on coverslips coated with Poly-L lysine (Invitrogen) in 24-well plates in Opti-MEM Reduced Serum Medium, GlutaMAX Supplement (Gibco). 24 h later, polymorphs were combined with Opti-MEM to a concentration of 2μM. Lipofectamine2000 (Invitrogen) was mixed with Opti-MEM (1:20 dilution) and incubated for 5 min. Lipofectamine mixture was then added 1:1 with α-Syn polymorphs and incubated for 20 min and then added to the cells to a final concentration of 100 nM. Labeled monomers of α-Syn were seeded as above with final concentration of 1μM. For seeding longer than 12 h, the media was exchanged for complete DMEM media. The cells were fixed in 4% PFA in PBS for 10 min. After washing 3x in PBS, the cells were mounted for imaging. For fluorescence lifetime imaging on Fibrils-Atto647 with no cells, Fibrils were incubated with lipofectamine as well and then seeded and fixed in the same manner as listed above.

MG132 (ThermoScientific) was dissolved in ethanol (EtOH) and chloroquine (Sigma) dissolved in H2O and were stored at −80 °C until use. For proteasomal inhibition experiments, cells were treated with MG132 at a working concentration of 5 μM for 4h before seeding with Fibrils for 12h. For autophagy inhibition experiments cells were treated with chloroquine at a working concentration of 40 μM for 12h before seeding with Fibrils for 12h. After fiber seeding for 12 h, the cells were fixed and then imaged.

For knock down of DNAJB1 and DNAJA2 treatment, ON-Target plus SMART Pool Human DNAJB1, ON-Target plus SMART Pool Human DNAJA2 and ON-Target plus Control Pool were purchased from ThermoScientific. Due to the long experimental timeframe, cells were seeded in Opti-MEM with 10%FBS. siRNA was diluted in siRNA buffer (Final concentration 20 nM) before mixing with Dharmafect for 20 min prior to being added to cells. After 48h knockdown, media was changed to Opti-MEM alone to seed with polymorphs for 24h as described above.

### Confocal Imaging

Cells were imaged with a Leica TCS SP8 STED 3X microscope (Leica Microsystem, Germany) equipped with 470-670nm white laser with a photomultipler tube and Leica Application Suite X (LASX) software using a HCX PL APO 63x/1.40 Oil CS2 objective. All further processing of acquired images was performed with ImageJ software(Schneider, Rasband, & Eliceiri, 2012). Manually quantification of cells and cells with fibrillar and spherical foci was done in FIJI (Schindelin et al., 2012) and the percentage of foci-containing cells was calculated.

### Fluorescence lifetime imaging microscopy

For FLIM imaging, Leica SP8 FALCON FLIM with a time-correlated single photon counting (TCSPC) module was used (Leica microsystem, Germany). A pulsed, white light laser provided excitation at 470-670nm and a repetition rate of 80 MHz. Images were acquired with 100X objective (HC PL APO 100x/1.40 STED White Oil) with zoom was applied up to 4-fold. Default FLIM settings for 514 nm and 633 nm excitation were applied for α-SynA53T-YFP and α-Syn-Atto647 polymorphs, respectively with two separate Hybrid GaAsp detectors in sequential mode (Leica microsystem, Germany). Data for FLIM were acquired until at least 1500 counts were collected in the brightest pixel of the image. Photons per laser pulse were kept under 0.5 to avoid photon pile up. TCSPC images were analyzed using FALCON FLIM (Leica) from which lifetime histograms and phasor plots were generated.

### SDS-Page and immunoblotting

Cells were centrifuged to pellet followed by lysis in lysis buffer (10mM Tris, 100mM NaCl, 0.2% Triton X-100, 10mM EDTA) on ice for 20min. The lysates were transferred into fresh Eppendorf tubes and centrifuged (1,000 g for 1 min at 4°C) in a tabletop centrifuge. The protein concentration was determined using protein assay dye reagent concentrate (Bio-Rad). Proteins were separated under denaturing conditions by SDS–PAGE and transferred onto a PVDF membrane (Carl Roth) by standard wet blotting protocols. Samples were probed with mouse monoclonal anti-HSP40/Hdj1 (2E1, Enzo) or monoclonal anti-DNAJA2 (ab147216, Abcam) primary antibody. Anti-GAPDH antibody (clone GAPDH-71.1, Sigma-Aldrich) was used as loading control. Alkaline phosphatase (AP)-conjugated anti-mouse IgG secondary antibodies (Vector Laboratories) were used for subsequent ECF-based detection (GE Healthcare).

### Statistical analysis

GraphPad Prism software was used to create graphs and to analyze the data. Data are presented as mean ± standard error of the mean (SEM). For each data set, p values and statistical test applied are indicated in the corresponding figure legend with the following significance levels: not significant (n.s.) p >0.05; *, p ≥0.05; **, p ≥0.01; and ***, p≥0.001.

## Acknowledgements

We thank Tracy Bellande for technical assistance and Holger Lorenz from the ZMBH Imaging Facility for his technical support and constructive comments on the manuscript. This work also benefited from the Microscopy platform of I2BC (CNRS UMR9198, Gif-sur-Yvette, France). We are also grateful to Marc Diamond for sharing his HEK293 α-SynA53T-YFP biosensor cell line. This is an EU Joint Programme - Neurodegenerative Disease Research (JPND) project (PROTEST-70). This project is supported through the following funding organizations under the aegis of JPND - www.jpnd.eu: France, Agence National de la Recherche (ANR, ANR-17-JPCD-0005-01 to R.M.); Germany, Bundesministerium für Bildung und Forschung (01ED1807B to C.N-K.). Funding was also provided by the Fondation pour la Recherche Medicale (contract DEQ. 20160334896), and France Parkinson Association.

## Conflict of interest

The authors declare no conflicts of interest.

## Contributions

Conceptualization, J.T. and C.N.-K.; Methodology, J.T., A.A., S.D.-A., R.M., and C.N.-K.; Investigation, J.T., S.D.-A.; Formal Analysis, J.T.; Resources, A.A., R.M.; Writing - Original Draft, J.T., C.N.-K.; Writing - Review and Editing, J.T., R.M., and C.N.-K.; Supervision, R.M. and C.N.-K.; Visualization, J.T.; Funding Acquisition, R.M., and C.N.-K.

---

**Supplementary Figure 1.**
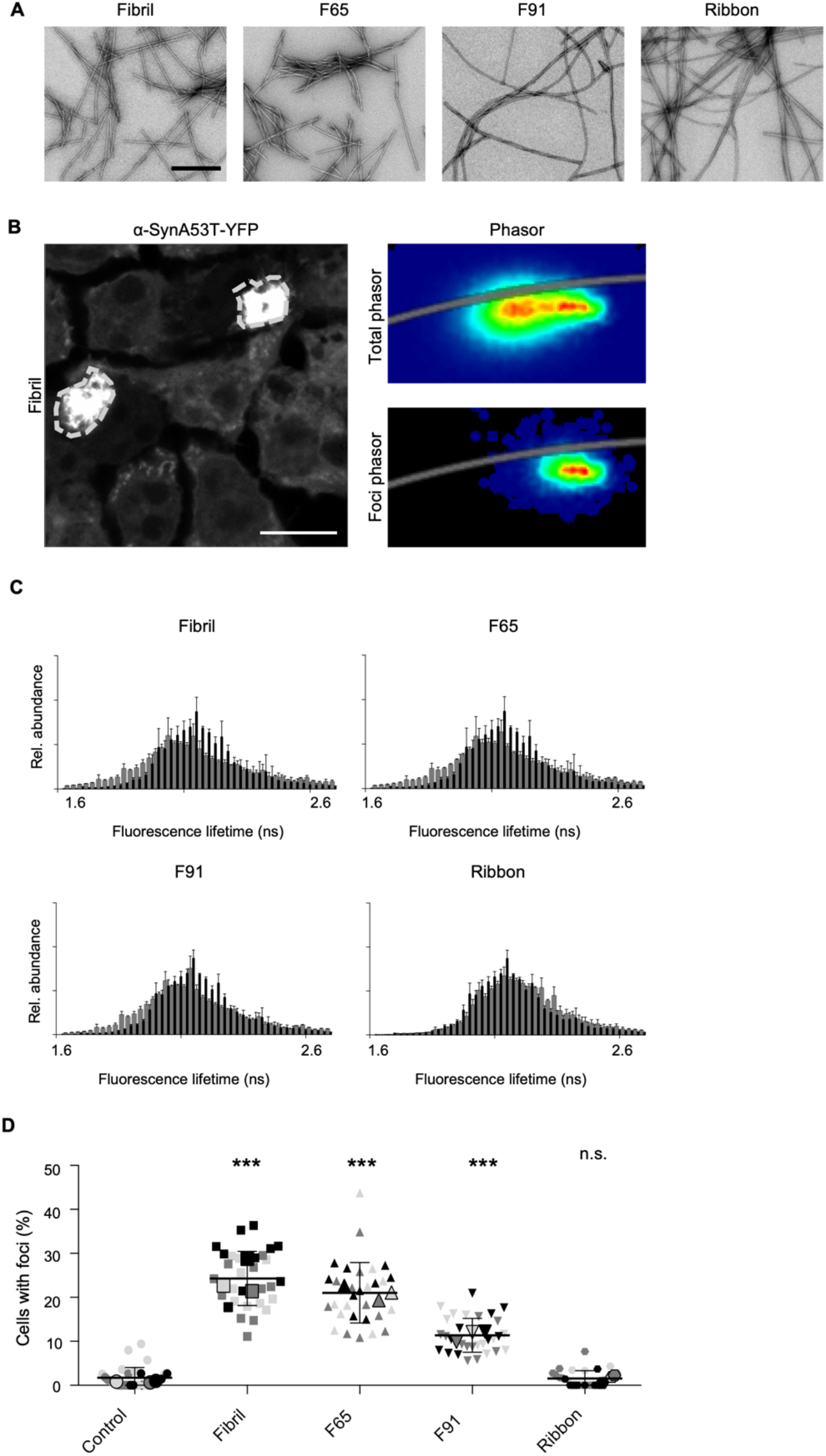
(A) Transmission electron microscopy (TEM) images of the different α-Syn polymorphs used throughout this study. Scale bar, 200 nm. (B) ROI drawn around foci in the fluorescence intensity image (dotted white line) induced by seeded aggregation show that foci contain α-SynA53T-YFP species with shorter lifetimes compared to the total phasor distribution. Scale bar, 10 μm. (C) Fluorescence lifetime histogram of α-SynA53T-YFP seeded with indicated α-Syn polymorphs, averaged over all acquired images (n = 25-30 for each polymorph, mean + SEM). Seeding with α-Syn polymorphs resulted in a small shoulder of shorter lifetimes in the overall distribution compared to the lifetime distribution of α-SynA53T-YFP in non-seeded cells (black). (D) Quantification of cells with visible foci. For each biological replicate (n = 3) a mean was calculated; these three means were then averaged (horizontal bars), and used to calculate the standard deviation (error bars), and p values. n.s. = not significant, ***p < 0.001. Statistical analysis was done using one-way ANOVA with Dunnett’s multiple comparison test.

**Supplementary Figure 2.**
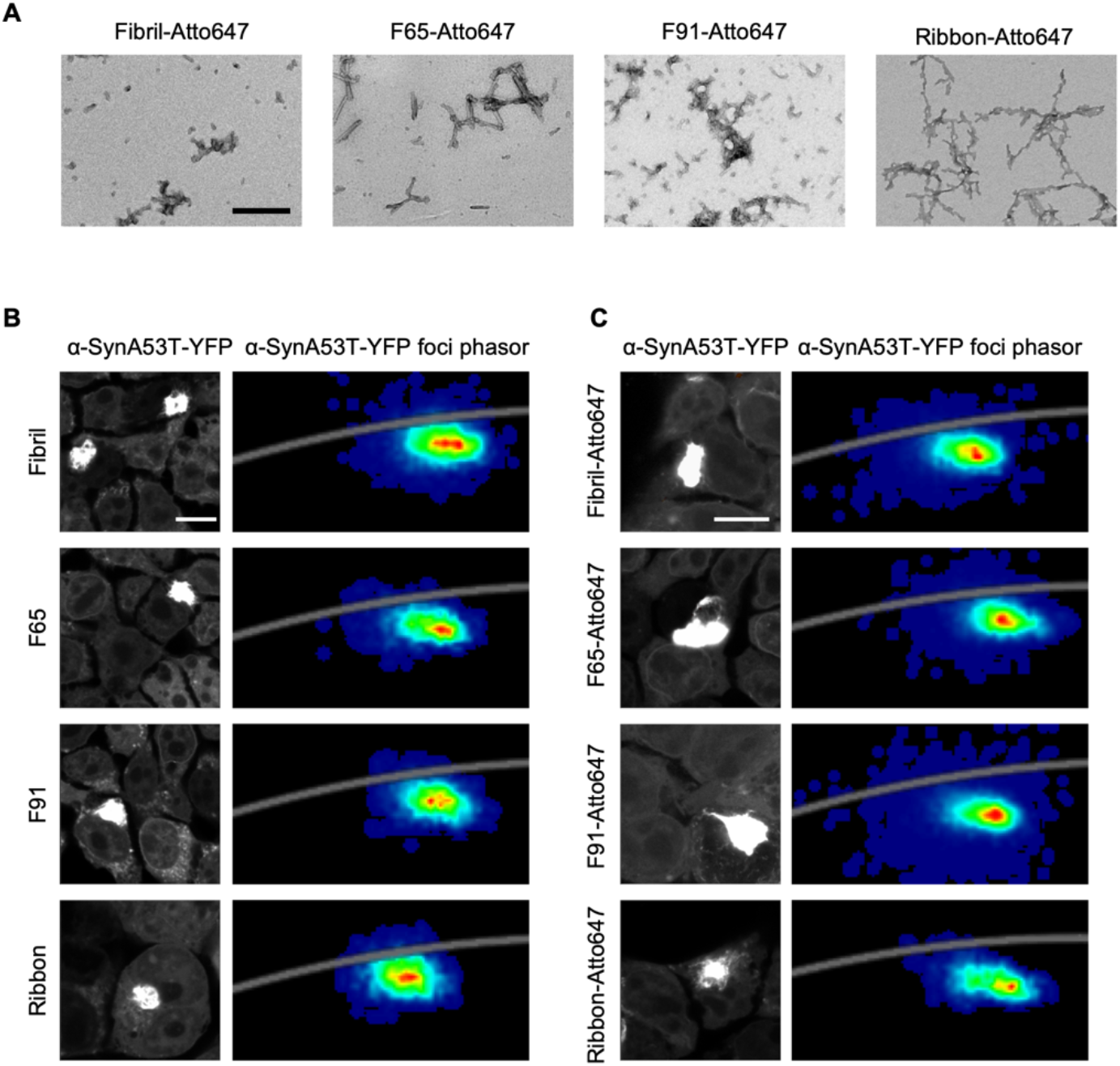
(A) Transmission electron microscopy (TEM) images of the different α-Syn polymorphs labeled with Atto647 used throughout this study. Scale bar, 200 nm. (B, C) Fluorescence intensity images of α-SynA53T-YFP seeded with either unlabeled or Atto647-labeled polymorphs α-Syn polymorphs. Corresponding phasor plots show the fluorescence lifetimes of the foci in α-SynA53T-YFP with seeded aggregation. Scale bar, 10 μm.

**Supplementary Figure 3.**
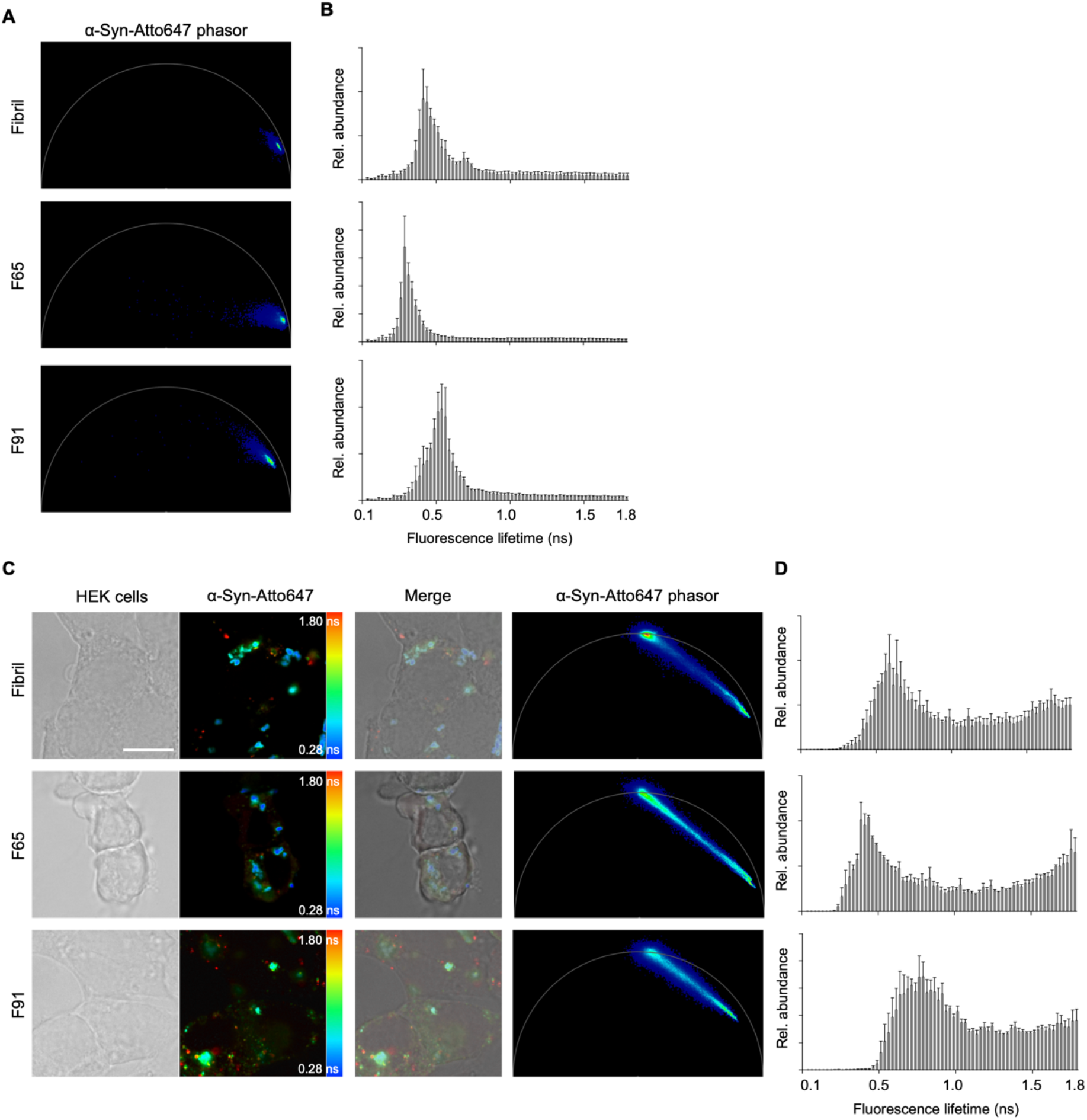
(A) Phasor plot of the fluorescence lifetime of α-Syn-Atto647 seeds when not added to cells (B) Histograms of α-Syn-Atto647 showing the relative abundance (rel. abundance) of pixels exhibiting a certain fluorescence lifetime (ranging from 0 to 1.8 ns), averaged over all acquired images (n = 15-20 for each polymorph, mean +SEM). (C) Transmitted light image of HEK293 cells and fluorescence intensity image of pseudo colored α-Syn-Atto647 seeds when added to cells with the merged image and the corresponding phasor plot of the fluorescence lifetime. α-Syn polymorphs retain their signature phasor fingerprint when added to HEK cells not expressing α-SynA53T-YFP. (D) Histograms of α-Syn-Atto647 seeds showing the relative abundance (rel. abundance) of pixels with a certain fluorescence lifetime (ranging from 0 to 1.8 ns), averaged over all acquired images (n = 15, mean + SEM). Scale bar, 10 μm.

**Supplementary Figure 4.**
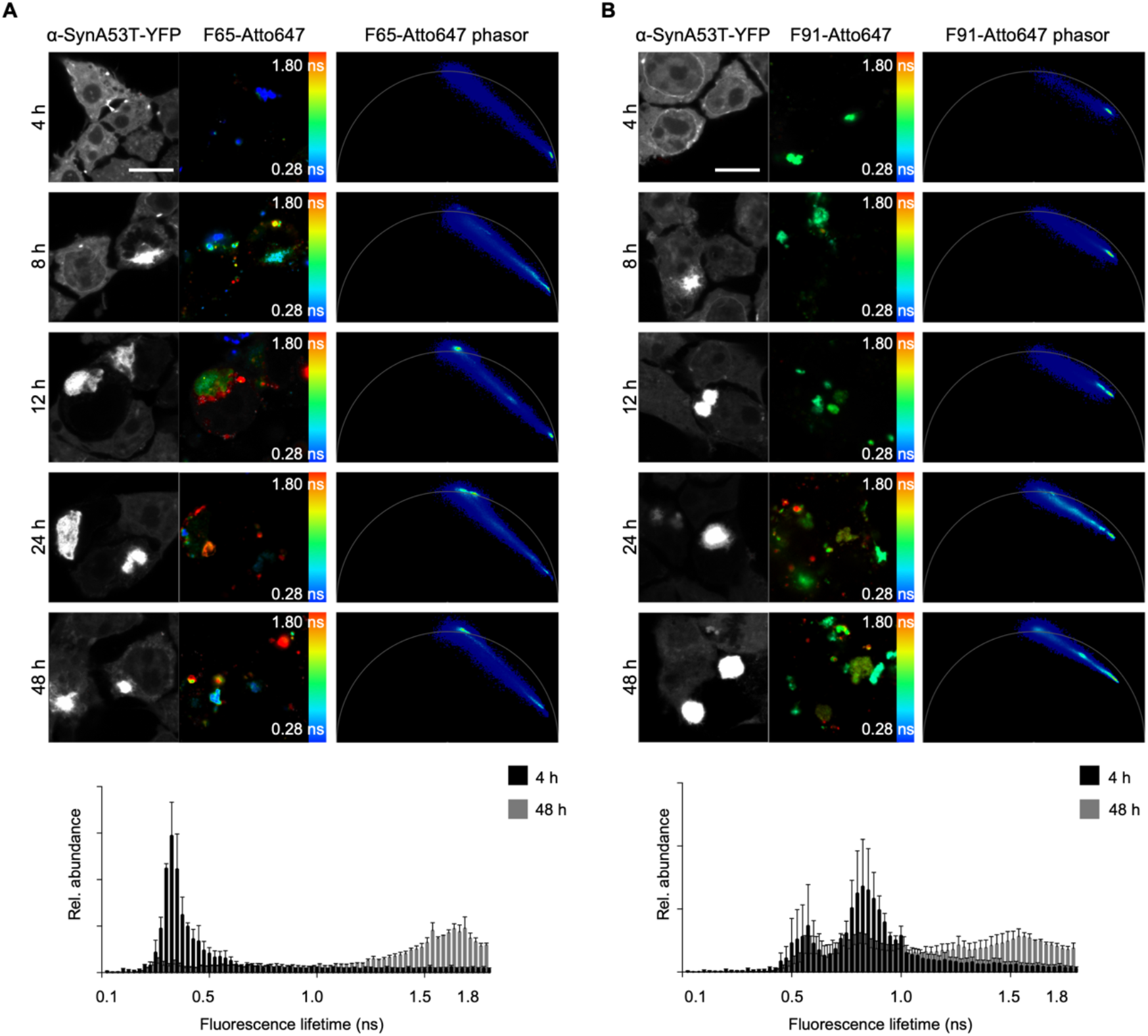
(A, B) Fluorescence intensity images of α-SynA53T-YFP and indicated α-Syn polymorphs labeled with Atto647 pseudo colored from blue (0.28 ns) to red (1.8 ns) to illustrate short (blue) and long (red) fluorescence lifetimes with the merged images and the corresponding phasor plots of the fluorescence lifetime of seeds at indicated timepoints after seeding. Histograms of α-Syn-Atto647 seeds showing the relative abundance (rel. abundance) of pixels with a certain fluorescence lifetime (ranging from 0 to 1.8 ns), at 4h and 48h, averaged over all acquired images (n = 15 for each timepoint, mean + SEM). F65 and F91 seeds accumulate species with longer fluorescence lifetimes in a time dependent manner. Scale bar, 10 μm.

**Supplementary Figure 5.**
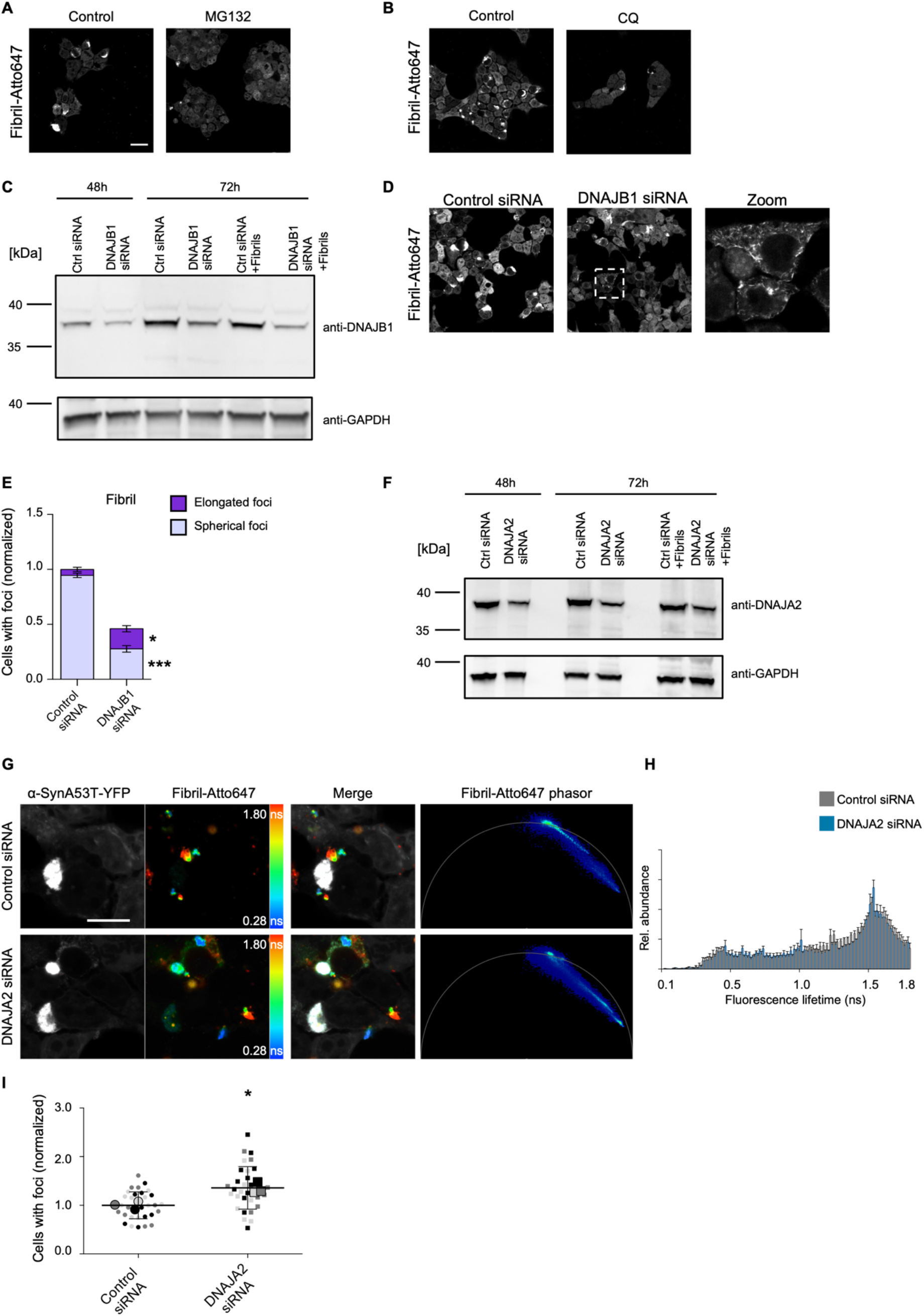
(A, B) Representative fluorescence intensity images of α-SynA53T-YFP cells seeded with Fibril-Atto647 and treated with MG132 or chloroquine (CQ). Scale bar, 20 μm. (C) Western blot to detect DNAJB1 levels when Fibril-Alex647 are added (48h) and after seeding for 24h (72h). Anti-GAPDH was used as a loading control. (D) Representative image of α-SynA53T-YFP cells seeded with Fibril-Atto647 treated with control or DNAJB1 siRNA. Scale bar, 20 μm. Zoom shows elongated foci seen in cells seeded with Fibril-Atto647 and treated with DNAJB1 siRNA (E) Quantification of cells with visible elongated and spherical foci upon DNJAB1 siRNA. *p < 0.05, ***p < 0.001. Statistical analysis was done using two-way ANOVA with Sidak’s multiple comparison test. (F) Western blot to detect DNAJA2 levels when Fibril-Atto647 seeds are added (48h) and after seeding for 24h (72h). Anti-GAPDH was used as a loading control. (G) Fluorescence intensity images of α-SynA53T-YFP and Fibril-Atto647 seeds pseudo colored from blue (0.28 ns) to red (1.8 ns) to indicate short (blue) and long (red) fluorescence lifetimes with the merged images and the corresponding phasor plots of the fluorescence lifetime of Fibril-Atto647 seeds upon treatment with control or DNAJA2 siRNA. Scale bar, 10 μm. (H) Histograms of Fibril-Atto647 showing the relative abundance (rel. abundance) of pixels with a certain fluorescence lifetime, averaged over all acquired images (n = 15 for each condition, mean + SEM). (I) Quantification of cells with visible foci after seeding with Fibril-Atto647. For each biological replicate (n = 3) the mean was calculated; these three means were then averaged (horizontal bars) and used to calculate standard deviation (error bars), and p values *p < 0.05. Statistical analysis was done using an unpaired t test.

**Supplementary Figure 6.**
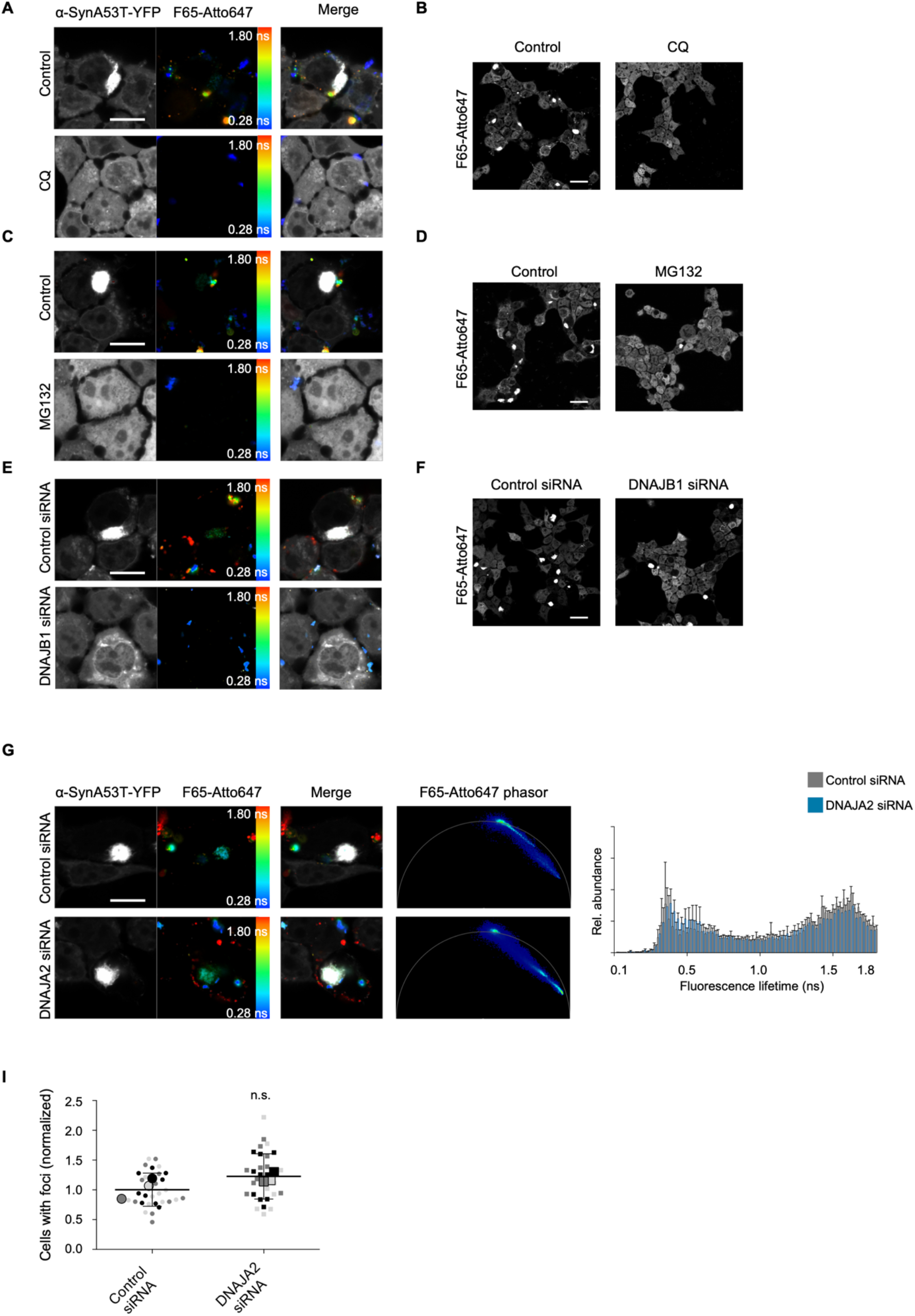
(A, C, E) Fluorescence intensity images of α-SynA53T-YFP and F65-Atto647 seeds pseudo colored from blue (0.28 ns) to red (1.8 ns) to illustrate short (blue) and long (red) fluorescence lifetimes with the merged images of the fluorescence lifetime of F65-Atto647 seeds upon treatment with chloroquine (CQ) (A), MG132 (C), or DNAJB1 siRNA (E), which correspond to the phasor plots in Figure 5. Scale bar, 10 μm. (B, D, F) Representative fluorescence intensity images of α-SynA53T-YFP cells seeded with F65-Atto647 and treated with chloroquine (CQ) (B), MG132 (D) or DNAJB1 siRNA (F). Scale bar, 20 μm. (G) Fluorescence intensity images of α-SynA53T-YFP and F65-Atto647 seeds pseudo colored and the corresponding phasor plots of the fluorescence lifetime of F65-Atto647 seeds upon treatment with control or DNAJA2 siRNA. Scale bar, 10 μm. (H) Histograms of F65-Atto647 showing the relative abundance (rel. abundance) of pixels with a certain fluorescence lifetime, averaged over all acquired images (n = 15 for each condition, mean + SEM). (I) Quantification of cells with visible foci after seeding with F65-Atto647. For each biological replicate (n = 3) the mean was calculated; these three means were then averaged (horizontal bars) and used to calculate standard deviation (error bars), and p values. n.s. = not significant. Statistical analysis was done using an unpaired t test.

**Supplementary Figure 7.**
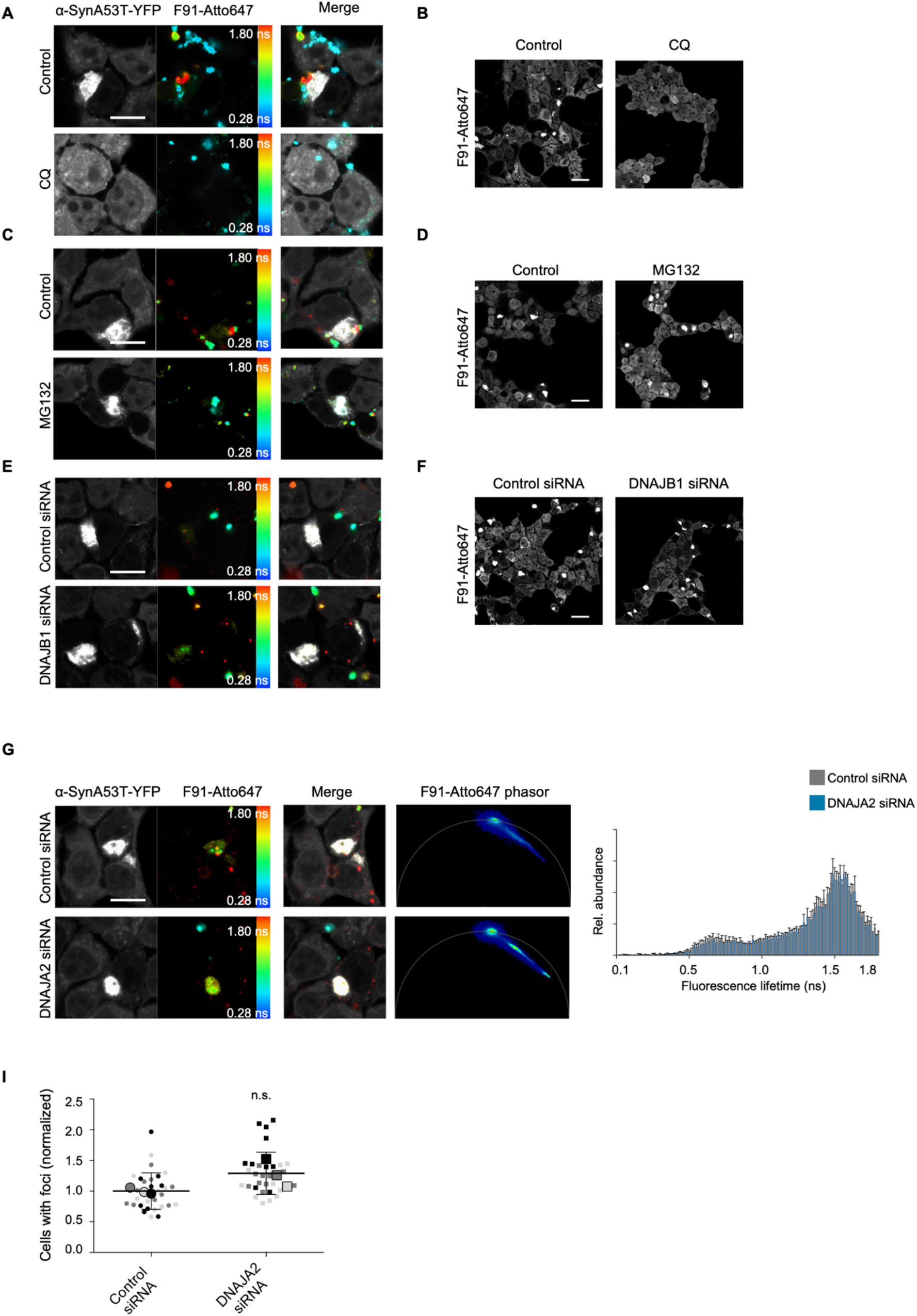
(A, C, E) Fluorescence intensity images of α-SynA53T-YFP and F91-Atto647 seeds pseudo colored from blue (0.28 ns) to red (1.8 ns) to illustrate short (blue) and long (red) fluorescence lifetimes with the merged images of the fluorescence lifetime of F91-Atto647 seeds upon treatment with chloroquine (CQ) (A), MG132 (C), or DNAJB1 siRNA (E), which correspond to phasor plots in Figure 5. Scale bar, 10 μm. (B, D, F) Representative fluorescence intensity images of α-SynA53T-YFP cells seeded with F91-Atto647 and treated with chloroquine (CQ) (B), MG132 (D) or DNAJB1 siRNA (F). Scale bar, 20 μm. (G) Fluorescence intensity images of α-SynA53T-YFP and F91-Atto647 seeds pseudo colored and the corresponding phasor plots of the fluorescence lifetime of F91-Atto647 seeds upon treatment with control or DNAJA2 siRNA. Scale bar, 10 μm. (H) Histograms of F91-Atto647 showing the relative abundance (rel. abundance) of pixels with a certain fluorescence lifetime, averaged over all acquired images (n = 15 for each condition, mean + SEM). (I) Quantification of cells with visible foci after seeding with F91-Atto647. For each biological replicate (n = 3) the mean was calculated; these three means were then averaged (horizontal bars) and used to calculate standard deviation (error bars), and p values. n.s. = not significant. Statistical analysis was done using an unpaired t test.

## Notes

### Competing Interest Statement

The authors have declared no competing interest.

